# When Task-Specific Learning Outperforms Transfer Learning: A Benchmark of Gene and Expression Encoding Strategies

**DOI:** 10.64898/2025.12.05.690830

**Authors:** Igor Sadalski

**Affiliations:** Somite.ai, Boston, MA, USA

## Abstract

Single-cell foundational models have emerged as a powerful tool for learning generalizable cellular representations from large-scale data. Most models in this domain use transformer backbones, which require careful engineering of gene and expression encoding strategies, yet there is no consensus on which encoding techniques are effective. While benchmarking efforts up to date have focused on evaluating downstream applications using already pretrained models, we take a fundamentally different approach: we isolate different encoding paradigms and systematically compare them by training models from scratch under controlled conditions. Moreover, we scale pretraining to 10 million cells across 100 diverse datasets, a tenfold increase compared to similar studies. Through empirical experiments, we find that contrary to common assumptions, pretrained embeddings from large protein models like ESM-2 consistently underperformed task-specific learned embeddings. Our work provides clear empirical guidance for model design decisions and establishes a systematic benchmark for evaluating encoding strategies in single-cell foundational models.

## 1 Introduction

A major driver in recent years has been to build an AI virtual cell, i.e., multi-scale, multi-modal neural network models that can represent and simulate cellular behaviour across diverse states (Bunne et al., 2024). Leading models promise to enable universal embeddings (Rosen et al., 2023), crossspecies transfer (Pearce et al., 2025), multi-task transfer learning for downstream applications (Cui et al., 2024; Theodoris et al., 2023), and batch correction (Wang et al., 2021). While foundational models can be generated for different omics data types (e.g., protein, transcriptomics, tissues), here we focus on transcriptomic data. In this domain, most of the models (e.g. Cui et al. (2024); Adduri et al. (2025); Pearce et al. (2025)) use a transformer (Vaswani et al., 2017) backbone. Since transcriptomic data inherently consists of two pieces of information, gene identities and their expressions, architects of these models are faced with a design choice on how to encode this information into embeddings that transformers can process.

Different encoding strategies embody distinct assumptions about what information is most useful. For genes, task-specific patterns versus biological priors, for expression values, continuous precision versus discrete robustness. Figure 2 provides a schematic overview of representative encoding paradigms. For gene encoding, some models learn embeddings de novo to capture dataset-specific co-expression relationships (Cui et al., 2024), while others leverage transfer learning from pretrained protein language models such as ESM-2 (Lin et al., 2023; Adduri et al., 2025) to incorporate evolutionary biological knowledge. For expression encoding, strategies range from discretizing into bins for computational efficiency (Cui et al., 2024; Gandhi et al., 2025) to using continuous or soft binning via MLPs (Pearce et al., 2025; Ho et al., 2024; Adduri et al., 2025) or applying binning based on logarithmic transformation (Wang et al., 2021).

**Figure 1.**
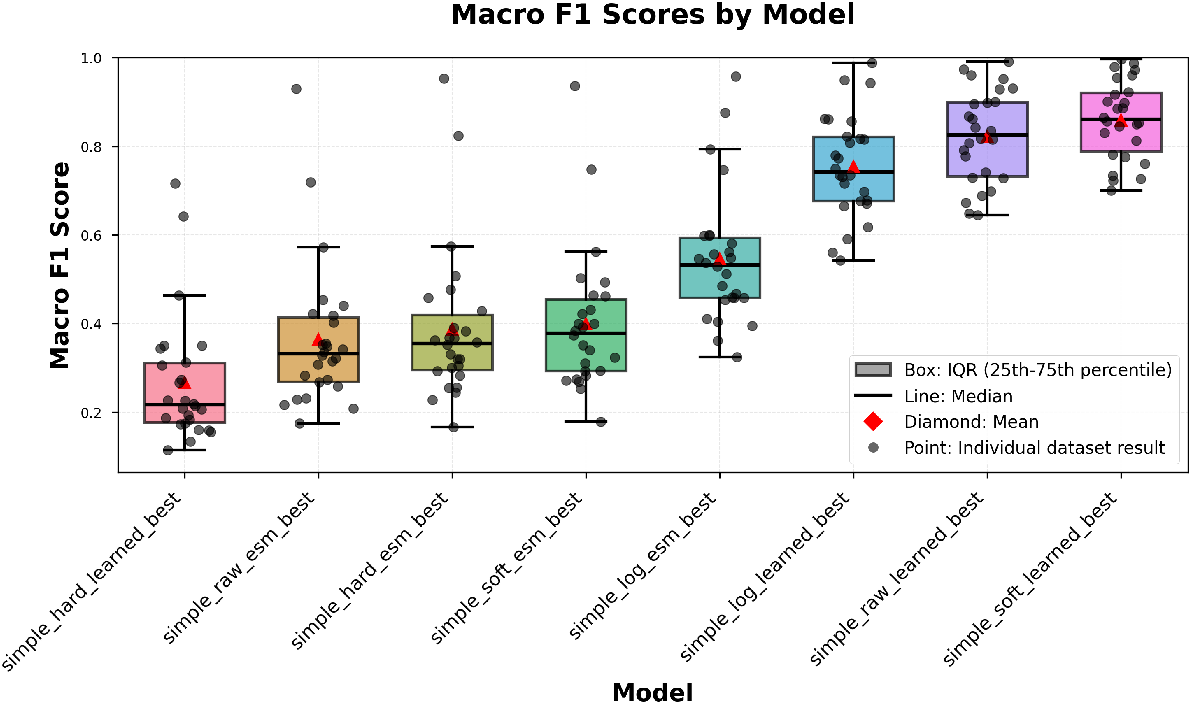
Boxplot of macro F1 score distribution by gene and expression encoding methods. Each point represents a macro F1 score for a kNN classifier trained on one of the evaluation datasets (Table 4) with a fixed trained transformer model.

**Figure 2.**
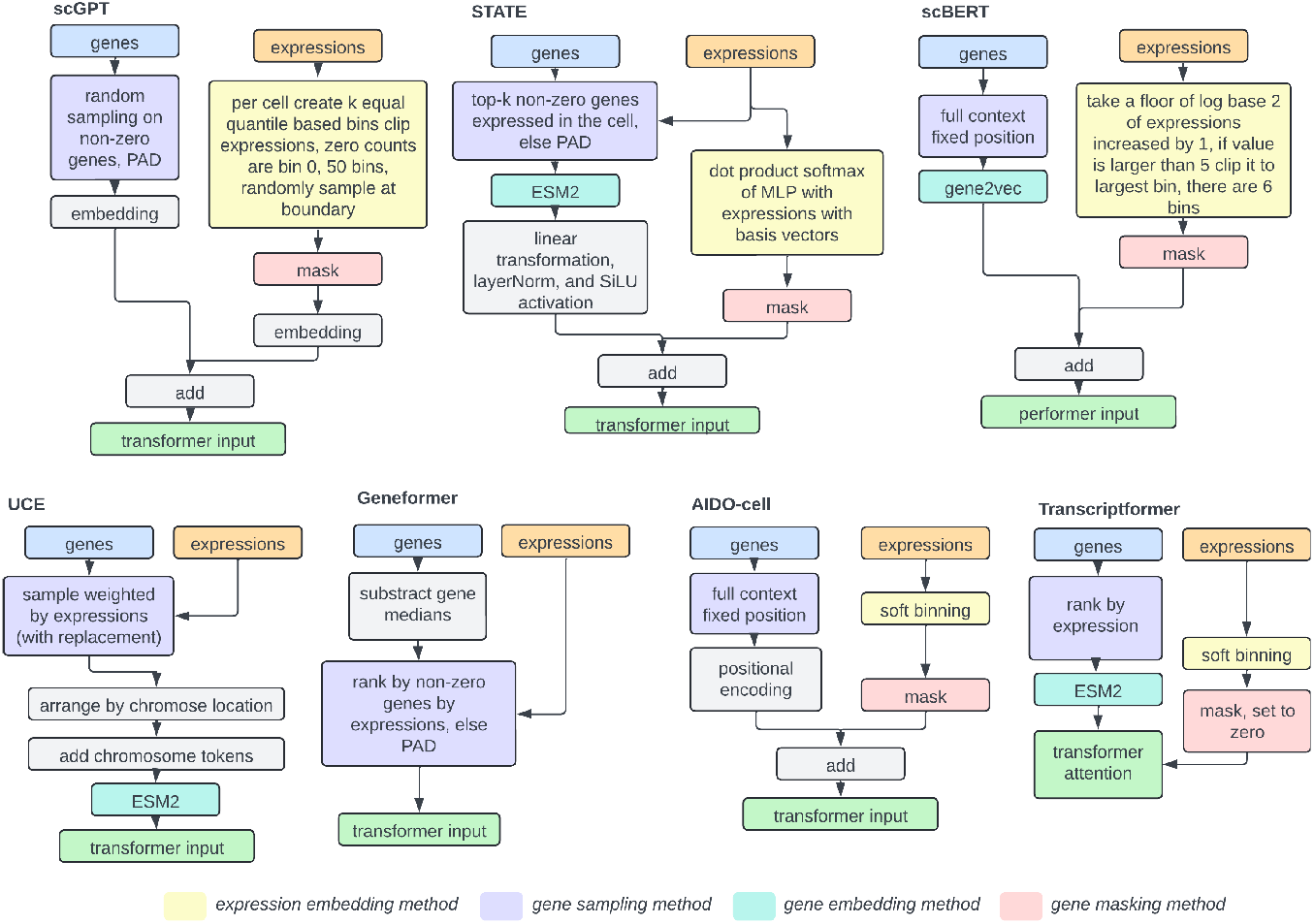
Schematic comparison of different gene and expression encoding methods across single cell foundational models (Cui et al., 2024; Adduri et al., 2025; Ho et al., 2024; Wang et al., 2021; Theodoris et al., 2023; Pearce et al., 2025; Rosen et al., 2023).

Limited scientific literature has addressed this problem. Most benchmarking efforts have focused on evaluating downstream applications, such as perturbation prediction (Ahlmann-Eltze et al., 2024; Wenteler et al., 2024), using already pretrained models. In this work, we focus more on specific architectural choices and pretrain our models from scratch. A similar approach was taken in HEIM-DALL (Haber et al., 2025), a modular tokenization framework, where authors tried to evaluate different encoding strategies. However, they pretrain their transformer model on only a few datasets with a total of 1 million cells. In comparison, regular transformer models in the field are trained on, e.g., 266 million cells (Gandhi et al., 2025) and hundreds of diverse datasets. Given improved transformer performance with data scale (Kaplan et al., 2020), larger pretraining can lead to different results.

We present a controlled benchmark that quantifies these methodologies using consistent model architecture and training procedures at a large scale. We overcome the limitations of previous work by scaling up the experiments tenfold to 10 million cells, increasing dataset diversity by using 100 different datasets, introducing new tokenization strategies such as log binning and raw embedding, and using a standardized benchmark (Tabula Sapiens v2) for evaluation.

The main contributions of this work are:

- **Tenfold increase in scale of the experiment** to 10 million cells and increasing diversity of the datasets by using 100 different datasets.
- **Evaluating new expression encoding strategies** like log binning and raw embedding.
- **Adding new standardized benchmark** (Tabula Sapiens v2) for evaluation.

## 2 Training Setup

We represent cells as bags of gene-expression pairs. Each cell *i* contains *M*_*i*_ genes with non-zero expression values, represented as:

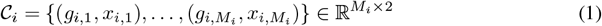

where *g*_*i,j*_ denotes the *j*-th gene identifier and *x*_*i,j*_ denotes its corresponding expression value in cell *i*.

Prior to encoding, we normalize expression values for each cell to a constant total *C* and apply log1p transformation:

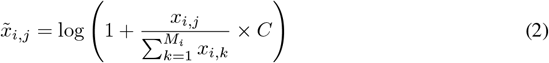

where *M*_*i*_ is the number of genes in cell *i, C* is a normalization constant (typically 10^4^ or 10^6^), and *k* indexes over all genes in cell *i*. This normalization step ensures consistent scaling across cells with varying sequencing depths.

Given computational constraints imposed by the context window size, we employ a sampling strategy to select a subset of genes for each cell. Let 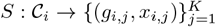 be a sampling function that takes cell *i* and returns *K* gene-expression pairs, where *K* = min(context window, *M*_*i*_). During training, for datasets where the number of non-zero expressed genes exceeds the context window, we randomly sample *K* genes from all non-zero expressed genes in each cell:

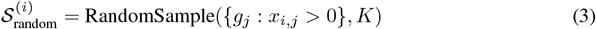

This approach ensures diverse gene representation across training examples while maintaining computational feasibility for large gene vocabularies.

The training objective is to predict masked expression values given the gene identities and unmasked expression context. To this end, we first compute embeddings for both gene identities and expression values. The gene embedding for each gene is obtained as:

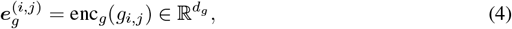

where enc_*g*_ is the gene encoding function (detailed in Section 3), *d*_*g*_ is the gene embedding dimension, and 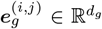 is the gene embedding vector. For expression values, we randomly mask a fraction *p*_mask_ of gene-expression pairs during training. The expression embedding is computed as:

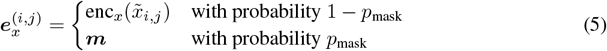

where enc_*x*_ is the expression encoding function (detailed in Section 4), 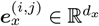 is the expression embedding vector, *p*_mask_ is the masking probability, and 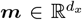 is a learnable mask token with dimension *d*_*x*_ matching the expression embedding dimension.

The combined gene-expression embedding for the *j*-th gene in cell *i* is obtained by summing the gene and expression embeddings:

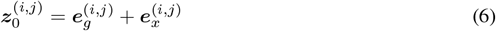

where 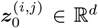 is the combined embedding for gene *j* in cell *i*, and *d* is the model dimension (equal to both *d*_*g*_ and *d*_*x*_). The combined embeddings for all *K* genes in cell *i* are concatenated together to form the input sequence for the model:

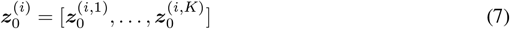

where 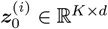 is the input sequence for cell *i* containing *K* gene embeddings. These combined embeddings are then passed through a transformer encoder. Grouping the gene-embeddings per cell and recursively applying the transformer layers, we obtain:

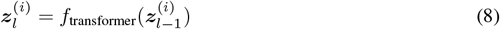

where *f*_transformer_ denotes a standard transformer encoder layer, *l* ∈ {1, …, *n*} indexes the layers, and *n* is the total number of transformer layers.

The predicted expression value for the *j*-th gene in cell *i* is decoded from the final transformer layer output using a simple MLP:

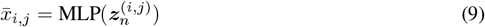

where 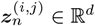 is the output embedding for gene *j* in cell *i* from the final transformer layer, and 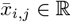 is the predicted expression value. We optimize the model using mean squared error loss for reconstruction of masked gene expression values (Wang et al., 2021; Adduri et al., 2025; Ho et al., 2024; Cui et al., 2024):

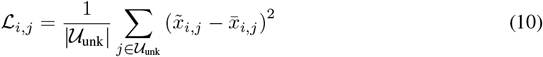

where ℒ_*i,j*_ is the loss for gene *j* in cell *i*, 𝒰_unk_ represents the set of masked gene indices, 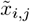 is the normalized true expression value, and 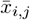 is the predicted expression value from Equation 9.

### 2.1 Evaluation

We evaluate model performance using cell type classification with a k-nearest neighbors (kNN) classifier applied to cell embeddings extracted from the final encoder layer (Pearce et al., 2025). For each cell, we extract its embedding vector from the transformer’s output, where the cell embedding is obtained by mean-pooling over all gene embeddings in the cell’s sequence.

The kNN classifier uses Euclidean distance to find the *k* = 10 nearest neighbors in the embedding space and assigns cell type labels based on majority voting among these neighbors. We set *k* = 10 as it provides a good balance between stability and sensitivity, and is a standard choice in the field. We also evaluated *k* = 5 and *k* = 15 and found similar relative rankings across configurations.

The primary metric is macro F1 score (Pearce et al., 2025), which provides balanced evaluation across all cell types by averaging per-class F1 scores:

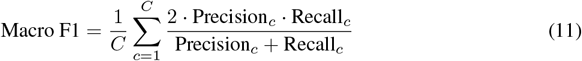

where *C* is the number of cell types, *c* indexes over cell types, and Precision_*c*_ and Recall_*c*_ are the precision and recall for cell type *c*, respectively.

## 3 Gene Encodings

**Learned encoding** is obtained by passing the gene identifier through a learned embedding table (Cui et al., 2024; Gandhi et al., 2025). This approach allows the model to generate task-specific representations for each gene, enabling it to capture dataset-specific relationships.

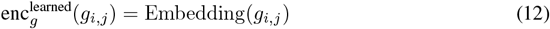

**ESM-2 encoding** uses precomputed embeddings retrieved from the ESM-2 (3B) model dictionary (Lin et al., 2023). STATE (Adduri et al., 2025) leverages this approach, using large pretrained protein language models to provide biologically-informed representations that are consistent across datasets and species, making it a strong choice for transfer learning and handling new or rare gene symbols. We project these embeddings through an MLP to match the model dimension.

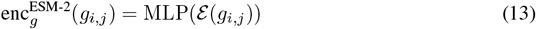

E where ℰ (*g*_*i,j*_) denotes the precomputed ESM-2 embedding for gene identifier *g*_*i,j*_, and MLP is a multi-layer perceptron that projects the embedding to the model dimension.

## 4 Expression Encodings

**Raw expression encoding** uses a simple MLP on normalized data. This approach preserves the full, continuous information from the expression measurement and is the most direct way to encode quantitative gene expression levels.

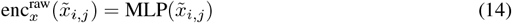

**Hard binning encoding** discretizes expression values into bins. This method groups expression levels into discrete intervals, trading off resolution for robustness and simplifying the input space (Cui et al., 2024; Gandhi et al., 2025).

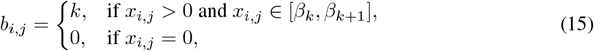

where *b*_*i,j*_ is the bin index for the expression value of gene *j* in cell *i, k* is the bin index, and *β*_*k*_ and *β*_*k*+1_ are the lower and upper boundaries of bin *k*, respectively. Expression embedding is obtained using an embedding layer:

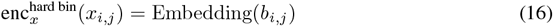

**Log binning encoding** compresses a wide dynamic range of expression values using a logarithmic transformation before discretizing. This approach can potentially mitigate the effects of outliers or skewed distributions. STATE (Adduri et al., 2025) uses this approach with ESM-2 embeddings, while scBERT (Wang et al., 2021) employs log binning with discrete tokenization.

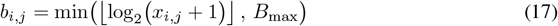

where *b*_*i,j*_ is the bin index for the expression value of gene *j* in cell *i*, and *B*_max_ is the maximum bin index. Bins are embedded using an embedding layer:

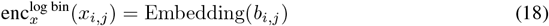

**Soft binning encoding** uses a softmax over potential bins to allow fractional/bin-weighted expression, which can capture uncertainty and subtle intensity differences between expression values, making the encoding differentiable and potentially more expressive (Ho et al., 2024; Hao et al., 2024; Pearce et al., 2025).

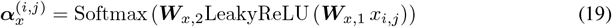

where 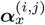 is a vector of bin weights for the expression value of gene *j* in cell *i*, ***W*** _*x*,1_ and ***W*** _*x*,2_ are learnable weight matrices, and LeakyReLU is the LeakyReLU activation function. The expression embedding is obtained by performing a soft lookup in the embedding table:

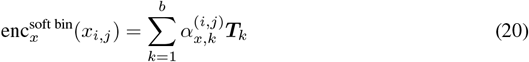

where *b* is the number of bins, 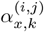 is the *k*-th element of 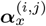 and ***T*** is the *k*-th embedding vector in the embedding table ***T***.

## 5 Data

### Training data

We train our models on 101 diverse single-cell RNA sequencing datasets comprising over 10 million cells, randomly selected from the curated collection used to train the Transcript-former model (Pearce et al., 2025). The datasets span diverse tissue types, experimental protocols, species, biological conditions, developmental stages, and disease states. See Appendix A for full details, including Table 3.

### Evaluation data

We evaluate all models on 26 tissue-specific datasets from the Tabula Sapiens v2 benchmark (Tabula Sapiens Consortium et al., 2022), comprising 548,977 cells across diverse human tissues. This benchmark represents a standard evaluation protocol in the field and has been used to evaluate other foundational models (Pearce et al., 2025). See Appendix A for full details, including Table 4.

## 6 Results

Table 1 summarizes kNN macro F1 score averages and standard deviations across all datasets that were used for evaluation. Learned gene embeddings consistently outperform ESM-2 embeddings across expression encoding methods, with the best configuration (soft binning + learned) achieving a mean macro F1 score of 0.860 ± 0.089 compared to 0.547 ± 0.152 for the best ESM-2 configuration (log binning + ESM-2), representing a 57% relative improvement. The performance gap between learned and ESM-2 embeddings is consistent across nearly all expression encoding methods, demonstrating that the choice of gene encoding has a larger impact than expression encoding. Figure 1 provides additional visualizations of the performance distributions across configurations.

**Table 1.** Performance comparison of different encoding configurations for gene and expression. Results show the mean kNN macro F1 score and standard deviation across 26 datasets used for validation.

| Expression Encoding | Gene Encoding | Mean Macro F1 (± std) |
| --- | --- | --- |
| **soft** | **learned** | **0.860 ± 0.089** |
| raw | learned | 0.823 ± 0.105 |
| log | learned | 0.755 ± 0.116 |
| log | esm | 0.547 ± 0.152 |
| soft | esm | 0.400 ± 0.160 |
| hard | esm | 0.388 ± 0.174 |
| raw | esm | 0.365 ± 0.164 |
| hard | learned | 0.268 ± 0.146 |

## 7 Discussion

Our benchmark reveals several important insights about encoding strategies for single-cell foundational models. The consistent outperformance of learned embeddings over ESM-2 embeddings challenges the assumption that transfer learning from protein language models provides superior representations for single-cell data.

### Why learned embeddings outperform ESM-2

We propose several explanations for this finding. First, learned embeddings can adapt to the specific statistical patterns in single-cell expression data, which differ substantially from protein sequence patterns. Second, the gene vocabulary in single-cell data includes many non-coding genes, pseudogenes, and gene isoforms that are not well-represented in protein language models. Third, learned embeddings capture co-expression relationships and gene-gene interactions that emerge from the training data, which are more relevant for downstream tasks than protein-level similarities.

### Why soft binning works best

Soft binning provides a balance between discrete and continuous representations. Unlike hard binning, which loses information at bin boundaries, soft binning allows the model to express uncertainty and capture subtle differences between expression values. Unlike raw continuous encoding, soft binning provides some regularization and structure that may help the model learn more robust representations. The differentiable nature of soft binning also enables end-to-end training, allowing the bin assignments to adapt to the task.

## 8 Limitations

Our study has several limitations that should be considered when interpreting the results. Our evaluation is limited to the Tabula Sapiens v2 benchmark (human tissues), cell type classification as the sole downstream task (as compared to work in (Author & Others, 2025)), and a single model architecture. The relative performance of encoding strategies may differ for other species, disease conditions, experimental protocols, or downstream tasks such as batch correction, perturbation prediction, or trajectory inference. Different architectures or hyperparameter tuning might also favour different encoding strategies.

Our training data consists of 101 datasets selected from a larger collection used for training single-cell foundational models. While the large scale (10M+ cells) should mitigate concerns about selection bias, the specific dataset choice may influence results. Additionally, our study focuses exclusively on encoding strategies and does not explore other important design choices such as attention mechanisms or training objectives. Future work should extend this benchmark to other architectural components and downstream tasks.

## 9 Conclusion

We present a large-scale systematic benchmark comparing gene and expression encoding strategies for single-cell foundational models. Our evaluation of 8 configurations trained on over 10 million cells demonstrates that (1) task-specific learned gene embeddings consistently outperform ESM-2 transfer learning, (2) soft binning provides the optimal expression encoding strategy, and (3) gene encoding choice has a greater impact than expression encoding on downstream performance. Future work should explore hybrid approaches, evaluate additional downstream tasks and species, and investigate alternative architectures. This benchmark contributes to the systematic evaluation of encoding strategies in single-cell modelling.

## Impact Statement

This paper presents a benchmark study that provides empirical guidance for designing effective foundational models in single-cell RNA sequencing analysis. Potential positive impacts include enabling more effective analysis tools for biomedical research and providing reproducible benchmarks for method comparison. Potential negative impacts are limited, as this is a methodological study focused on model architecture rather than direct applications.

## A Additional Details

### A.1 Reproducibility

Training datasets are from publicly available sources (Pearce et al., 2025). Evaluation datasets from Tabula Sapiens v2 are available from the official repository (Tabula Sapiens Consortium et al., 2022).

### A.2 Hyperparameters

Table 2 summarizes the hyperparameters used for training all models in this study.

**Table 2.** Hyperparameters used for model training.

| Hyperparameter | Value |
| --- | --- |
| *Model Architecture* |  |
| Model dimension ($d_{\text{model}}$) | 512 |
| Number of transformer blocks ($n_{\text{blocks}}$) | 12 |
| Number of attention heads ($n_{\text{head}}$) | 8 |
| Feed-forward dimension ($d_{\text{hid}}$) | 1024 |
| Dropout | 0.1 |
| Context window | 1024 |
| *Training* |  |
| Batch size | 512 |
| Gradient accumulation steps | 32 |
| Effective batch size | 16,384 |
| Epochs | 3 |
| Learning rate | $3 \times 10^{-4}$ |
| Weight decay | $10^{-5}$ |
| Optimizer betas | [0.9, 0.95] |
| Max gradient norm | 1.0 |
| Scheduler | Cosine annealing |
| Warmup steps | 2000 |
| Early stopping patience | 100 |
| *Data* |  |
| Validation ratio | 0.001 |
| Test ratio | 0.001 |
| Masking probability ($p_{\text{mask}}$) | 0.5 |
| *Expression Encoding* |  |
| Raw hidden dimension | 512 |
| Hard binning bins | 50 |
| Soft binning bins | 20 |
| Soft binning hidden dimension | 512 |
| Log binning max bins | 10 |
| *Other* |  |
| Random seed | 777 |
| Number of data workers | 8 |

### A.3 Training Datasets

Table 3 lists all 101 training datasets used in this study, including the dataset identifier and number of cells for each dataset. The total number of cells across all training datasets is 10,010,835.

**Table 3:**
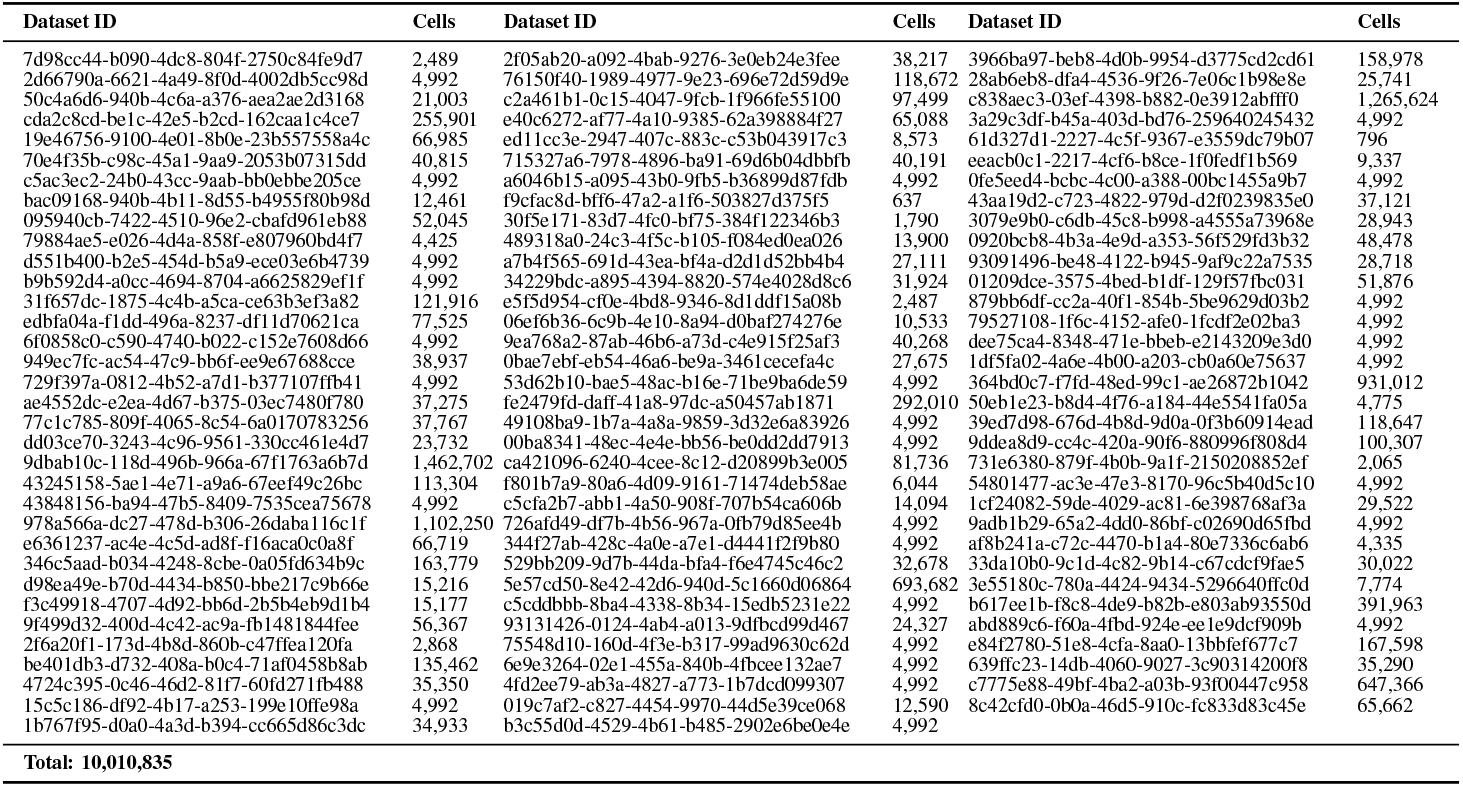
Training datasets used for model pretraining. Dataset IDs are UUIDs from the curated dataset collection.

### A.4 Evaluation Datasets

Table 4 lists all 26 evaluation datasets from the Tabula Sapiens v2 benchmark, including the tissue type and number of cells for each dataset.

**Table 4:** Distribution of datapoints in evaluation datasets from Tabula Sapiens v2 benchmark (Tabula Sapiens Consortium et al., 2022).

| Tissue | Number of Cells | Tissue | Number of Cells |
| --- | --- | --- | --- |
| Bladder | 36,715 | Blood | 17,802 |
| Bone Marrow | 8,045 | Ear | 3,055 |
| Eye | 21,121 | Fat | 71,021 |
| Heart | 13,344 | Large Intestine | 15,231 |
| Liver | 10,002 | Lung | 11,716 |
| Lymph Node | 72,290 | Mammary | 18,539 |
| Muscle | 13,035 | Ovary | 48,942 |
| Prostate | 2,044 | Salivary Gland | 11,561 |
| Skin | 6,486 | Small Intestine | 25,834 |
| Spleen | 30,690 | Stomach | 33,064 |
| Testis | 7,513 | Thymus | 9,933 |
| Tongue | 21,499 | Trachea | 5,346 |
| Uterus | 10,580 | Vasculature | 23,569 |
| **Total** |  |  | **548,977** |

## Notes

### Competing Interest Statement

The authors have declared no competing interest.

